# INFLUENCE OF RAT CENTRAL THALAMIC NEURONS ON FORAGING BEHAVIOR IN A HAZARDOUS ENVIRONMENT

**DOI:** 10.1101/2021.12.31.474632

**Authors:** Mohammad M. Herzallah, Alon Amir, Denis Paré

**Author notes:** <u>Correspondence should be sent to</u>: Denis Paré, Center for Molecular and Behavioral Neuroscience, Rutgers University - Newark, 197 University Avenue, Newark, NJ 07102.

## Abstract

The basolateral amygdala (BL) is a major regulator of foraging behavior. Following BL inactivation, rats become indifferent to predators. However, at odds with the view that the amygdala detects threats and generate defensive behaviors, most BL neurons have *reduced* firing rates during foraging and at proximity of the predator. In search of the signals determining this unexpected activity pattern, this study considered the contribution of the central medial thalamic nucleus (CMT), which sends a strong projection to BL, mostly targeting its principal neurons. Inactivation of CMT or BL with muscimol abolished the rats’ normally cautious behavior in the foraging task. Moreover, unit recordings revealed that CMT neurons showed large but heterogeneous activity changes during the foraging task, with many neurons decreasing or increasing their discharge rates, with a modest bias for the latter. A generalized linear model revealed that CMT neurons encode many of the same task variables as principal BL cells. However, the nature (inhibitory vs. excitatory) and relative magnitude of the activity modulations seen in CMT neurons differed markedly from those of principal BL cells but were very similar to those of fast-spiking BL interneurons. Together, these findings suggest that, during the foraging task, CMT inputs fire some principal BL neurons, recruiting feedback interneurons in BL, resulting in the widespread inhibition of principal BL cells.

## INTRODUCTION

In the wild, rodents face a perpetual dilemma: to stay in sheltered areas where resources are scarce or to explore open areas where they might find needed resources but also risk being detected by a predator. Biasing this life-and-death calculus are various factors such as current energetic needs, experience with the environment, and the palatability of potential resources (Mobbs et al. 2018). While multiple cortical (Rushworth and Behrens, 2008; Addicott et al., 2017; Scholl and Flugge, 2018) and subcortical sites (Choi and Kim, 2010; Canteras et al., 2012) determine the overall impact of these factors on foraging behavior, the basolateral complex of the amygdala (BLA) appears to exert a particularly profound influence.

Indeed, lesion and inactivation of the BLA drastically reduce the defensive behaviors normally elicited by predators, their odor, or associated contextual cues (Vazdajarnova et al., 2001; Takahashi et al., 2007; Choi and Kim, 2010; Martinez et al., 2011; Bindi et al., 2018). For instance, in a semi-naturalistic foraging task where rats are confronted with a mechanical predator when they venture out of their nest to forage, rats normally show extremely cautious behaviors including prolonged periods of hesitation at the edge of their nest and rapid escape when charged by the predator (Choi and Kim, 2010; Amir et al., 2015). In contrast, following BLA inactivation, rats quickly leave their nest and seem indifferent to the predator (Choi and Kim, 2010).

Unexpectedly however, unit recordings in the same task revealed that very few lateral (LA) and basolateral (BL) amygdala cells are responsive to the predator and that many of these rare neurons are also excited by non-threatening stimuli (Amir et al., 2015, 2019a; Kim et al., 2018). In fact, opposite to expectations, most principal BL neurons show *reduced* firing rates when rats initiate foraging and become nearly silent close to the predator (Amir et al., 2015, 2019a). Yet, when rats hesitate at the edge of their nest, BL cells fire at higher rates if rats eventually abort than start foraging or in the presence vs. absence of the predator (Amir et al., 2015, 2019a). Together, these results suggest a counterintuitive interpretation for the effects of BL inactivation in the foraging task: BL inactivation does not reduce defensive behaviors because it abolishes threat signaling by BL neurons but because it reproduces the firing suppression that normally develops when rats initiate foraging.

At present, the origin and nature of the signals responsible for this unexpected pattern of activity are unknown. Here, we examined the possibility that the central medial thalamic nucleus (CMT), which contributes very strong projections to BL (Vertes et al., 2015; Amir et al., 2019b), is involved. To this end, we compared the effect of inactivating CMT and neighboring sites on behavior in the foraging task. Moreover, we recorded unit activity in the CMT of rats performing the foraging task and compared their activity profile to that of BL neurons we recorded previously (Amir et al., 2015).

## MATERIAL AND METHODS

To test the hypothesis that CMT regulates behavior in the foraging task, we performed two sets of experiments. In the first, we compared the effects of inactivating BL, CMT, and neighboring thalamic sites on foraging behavior. In the second, we recorded CMT unit activity while rats performed the foraging task. These two experiments are described in turn below. All procedures were approved by the Institutional Animal Care and Use Committee of Rutgers University, in compliance with NIH’s Guide for the Care and Use of Laboratory Animals.

### Subjects

We used Sprague Dawley rats (Charles River, Fairfield, NJ; male; initial bodyweight of 250-275 g) housed individually and with *ad libitum* access to food and water. Rats were kept on a 12-hour light-dark cycle (lights off at 19:00 h) and experiments were conducted during the light phase of the cycle. During the period devoted to the foraging task, rats had restricted access to food to ensure proper motivation. During this period, their bodyweight was maintained at 90% of normal age-matched values.

### Behavioral experiment

#### Surgery

Figure 1A summarizes the experimental timeline of the behavioral experiments. Under aseptic conditions and deep isoflurane anesthesia, rats were placed in a stereotaxic apparatus with non-puncture ear bars (Kopf, Tujunga, CA). Rats received atropine sulfate (0.05 mg/kg, i.m.) to facilitate breathing and their scalp was injected with a local anesthetic (bupivacaine, 0.1-0.3 mL of a 0.125% solution, s.c.) in the area to be incised. Ten minutes later, we made a 0.5 cm scalp incision and a craniotomy above the brain region of interest. Under stereotaxic guidance, rats were implanted with guide cannulae (0.48 mm o.d., 0.32 mm i.d.; World Precision Instruments, Sarasota, FL) aimed at CMT (only one cannula) or BLA (one cannula per hemisphere). The cannulae were attached to the skull with dental cement and anchoring screws. They contained a dummy cannula to prevent blocking. At the conclusion of the surgery, rats were administered an analgesic with a long half-life (ketoprofen, 2 mg/kg, s.c., daily for three days). They were allowed a 7-day recovery period during which they were habituated to handling daily. We used the following stereotaxic coordinates (in mm relative to bregma): BLA (AP −2.4; ML ±5.00, DV −8.00; Angle from midsagittal line: 0°); CMT (AP −2.28, ML 0.4; DV −7.40, 20° mediolateral angle).

**Figure 1.**
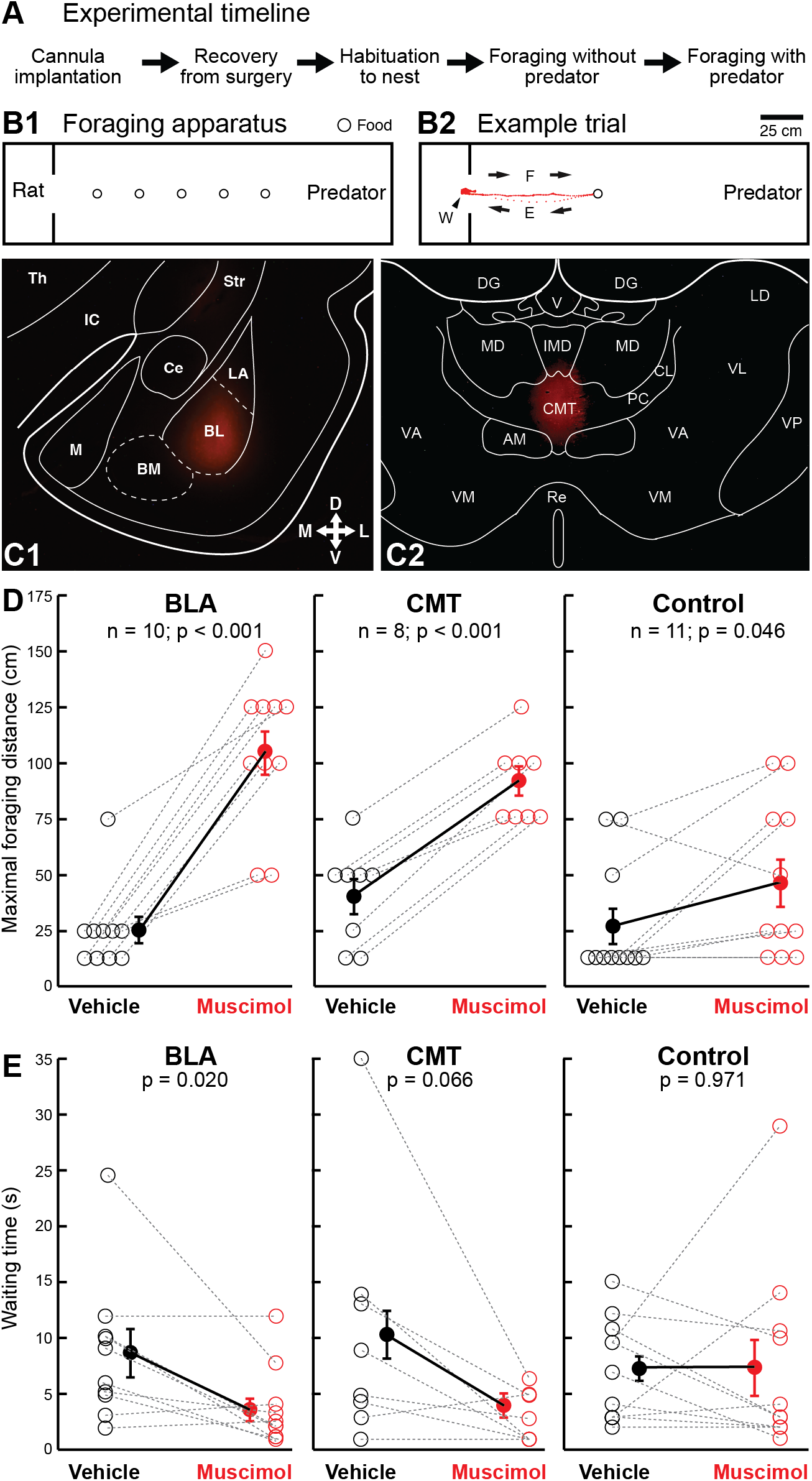
Effect of CMT inactivation on foraging behavior. (**A**) Experimental timeline. (**B1**) Top view on the behavioral apparatus, which was comprised of a small, dimly lit nesting area (left) and a brightly lit and much longer foraging arena (right). The two compartments were separated by a sliding door whose opening marked the start of a trial. On each trial, one food pellet was placed at various distances (circles) from the nest. (**B2**) Example trial. Red dots indicate rat position from the waiting phase (W), through the foraging (F) and escape (E) phases. Distance between dots is proportional to rat’s speed. (**C**) Examples of fluorescent muscimol infusion sites in the BLA (**C1**) and CMT (**C2**). (**D**) Maximal foraging distance or (**E**) waiting time when vehicle (black) or muscimol (red) was infused in the BLA (left), CMT (middle), or control sites (right). Empty symbols: individual subjects. Filled symbols, group averages ± SEM.

#### Foraging apparatus

The foraging apparatus (**Fig. 1B1**) was rectangular in shape and made of black polycarbonate. It had walls 60 cm high, and was divided in two compartments, both 60 cm wide. The first compartment was a small, dimly-lit nest (30 cm in length; 10 Lux) with a water bottle. The second was an elongated and brightly lit foraging arena (245 cm in length; 200 Lux). The two compartments were divided by a sliding door.

#### Behavioral procedures

Habituation (Days 1-2): Rats were first habituated to the nesting area (3.5 h/day for two days). During these sessions, the gateway to the nesting area was closed and rats could consume up to 6 g of food.

Foraging without the predator (Days 3-9): After habituation to the nest, rats underwent seven consecutive days of foraging training without the predator. After 60-90 s in the nest (no food), the door was opened and the rat was allowed to explore the foraging arena and search for a food pellet. On the first trial, the food pellet was placed 25 cm from the nest. After each successful trial, the distance was increased in steps of 25 cm up to 75 cm. Upon successful retrieval of the food pellet and reentry into the nest, the door was closed. One minute after the animal finished consuming the pellet, the gateway was reopened and another trial began. Each day, rats were required to complete at least three successful trials.

Foraging with predator (Days 10-11): During the last two days of the experiment, on each trial, we placed a robotic predator at the end of the foraging arena opposite to the nest, but facing it. The predator (Mindstorms, LEGO Systems, Billund, Denmark) was 14 cm tall, 17 cm wide, and 34 cm long. Each time rats approached within ~25 cm of the food pellet, the predator surged forward (80 cm at 60 cm/s), snapped its jaw repeatedly (~9 times), and returned to its original position. On the first trial of each experimental day, a food pellet was placed 75 cm from the nest. If the rat successfully retrieved the pellet, the distance was increased in steps of 25 cm until the rat failed. When the rats failed at 75 cm, the distance was decreased in steps of 25 cm until the rat retrieved the pellet. Trials were separated by 1-2 min intervals during which rats were in the nest with the door closed.

#### Drug Infusions

On Days 10-11, we performed infusions (0.1 μL/hemisphere) of artificial cerebrospinal fluid (aCSF, pH ~7.4) or fluorescent muscimol (0.9 nanomoles/hemisphere) in BLA (bilaterally) or CMT (only one infusion). The dummy cannula was removed. Both drugs were dissolved in aCSF and obtained from Sigma-Aldrich (St. Louis, MO). The composition of the aCSF (in mM) was 126 NaCl, 2.5 KCl, 1 MgCl_2_, 26 NaHCO_3_, 1.25 NaH_2_PO_4_, 2 CaCl_2_, 10 glucose, with pH 7.2, and 300 mOsm. The solution was backloaded in infusion cannulas (0.3 mm o.d.; 0.16 mm i.d.) using polyethylene tubing connected to 10 μL micro-syringes and infused at a rate of 0.1 μL/min using a microdialysis pump (CMA402, Harvard Apparatus, Holliston, MA). The infusion cannulae remained in place for ~60 s after the infusion. For inserting the infusion cannulae into the guide cannulae, rats were habituated to gentle restraint for 10-15 min daily for 3-5 d prior to the first infusion. At the end of this habituation period, rats could sit quietly on the investigator’s lap and groom for 10-12 min with minimal restraint. vehicle or muscimol infusions were performed on different days, 24 hours apart. To minimize the potentially confounding influence of habituation to the predator, the order of the infusions was counterbalanced in each group.

#### Analysis of behavior

Behavior was recorded by an overhead video camera at a frame rate of 29.97 Hz. A MATLAB script determined the rats’ position and velocity by taking advantage of the shifting distribution of light intensity across frames. Moreover, we analyzed the video files frame-by-frame to identify when rats started waiting at the door threshold (operationally defined as when their snout extended passed the door into the foraging arena), when they started foraging (operationally defined as the last frame of stillness prior to moving entirely out of the nest), retrieved a food pellet, escaped, and retreated into the nest (**Fig. 1B2**). We also noted whether rats succeeded or failed each trial. The start of the “escape” phase was defined as when rats, after approaching the food pellet, abruptly turned around to run back to the nest. This behavior was observed whether the predator was present or not and whether the trial was successful or not.

#### Histology

One day after completion of the behavioral tests, rats received a bilateral infusion of fluorescent muscimol, as described above. Two hours later, they were overdosed with isoflurane and perfused transcardially with 0.9% saline followed by 4% paraformaldehyde. Their brains were then stored in 4% paraformaldehyde overnight. A tissue block containing the region of interest was then sectioned in the coronal plane using a vibrating microtome (80 μm sections). The sections were mounted on gelatin-coated slides, air dried for two days in obscurity, cover-slipped with DPX-mounting solution (Sigma-Aldrich, St. Louis, MO), and examined with a fluorescence microscope (Nikon Eclipse E800, Melville, NY).

### Electrophysiological experiment

#### Surgical procedures

Surgical techniques were identical to those described above with the following exceptions. Under stereotaxic guidance, we aimed silicon probes (NeuroNexus, Ann Arbor, MI) to CMT. Silicon probes consisted four shanks (intershank distance of 200 μm), each with eight recording leads (de-insulated area of 144 μm^2^) separated by ~20 μm dorsoventrally. They were attached to Buzsaki-style microdrives (Vandecasteele et al., 2012) such that they could be lowered between recording sessions. Rats were allowed 2–3 weeks to recover from the surgery.

#### Foraging task

The foraging apparatus was identical to that used in the first experiment. So were the training stages and behavioral scoring procedures. However, after rats were trained on the foraging task in the absence of the predator (days 3-4), on each recording day (days 5-7), we conducted alternating trial blocks with (*n* = 10–20) or without (*n* = 10–15) the predator, for a total of 100–120 trials per day.

#### Histology

At the conclusions of the experiments, while under deep isoflurane anesthesia, electrolytic marking lesions were made on the most dorsal or ventral recording leads, alternating between shanks (10 μA for 16 s), so that lesions marking different shanks could be distinguished. Rats were then perfused-fixed trans-cardially, their brains extracted and sectioned. Sections were then counterstained with a 1% thionine solution. All neurons included in this study were histologically determined to have been recorded in CMT.

#### Electrophysiological procedures

The data was sampled at 25 kHz and stored on a hard drive. The data was high-pass filtered (cutoff 300 Hz), followed by a median filter (window of 1.1 ms). To extract action potentials, a threshold was applied to the filtered data. Using principal component analysis on the spike waveforms, the first three components were clustered using KlustaKwik (http://klustakwik.sourceforge.net/) and the resulting clusters of spikes were improved using Klusters (Hazan et al., 2006). To separate the clusters, we computed auto-correlograms and cross-correlograms for potential merging or additional splitting. For units to be considered for further analyses, autocorrelograms had to exhibit a refractory period of ≥2 ms. Cross-correlograms with a refractory period were merged as this implied that the same neuron was shared between clusters. Cells with unstable action potential waveforms were excluded.

#### Generalized Linear Model (GLM)

We fit the spiking activity of individual units using a regularized regression, group least absolute shrinkage and selection operator (Lasso) with Poisson distribution (grpreg version 3.2-0 R package; Breheny and Huang 2015) with sevenfold cross-validation, identical to the GLM used by Amir et al. (2019a). Spiking was time-binned (66.7 ms bins, which corresponds to two video frames). Trials included four task phases: door opening when rats were in the nest, waiting at the door, foraging, and escape. Task variables considered in the GLM were door opening, waiting, foraging, escape, food retrieval, predator activation, nest reentry, speed, distance from nest, distance from food, trial type (without or with predator), and prior trial outcome (failure or success). The input was coded at the initiation of the variable (door opening, predator activation), throughout the length of the variable (waiting, foraging, escape, food retrieval, nest reentry), or throughout the length of the trial (trial type, prior trial outcome, speed, distance from nest, distance from food). Variables were either coded as dummy (waiting, foraging, escape, trial type, prior trial) or convolved with a group of 14 basis functions defined by log-timescaled-raised cosine bumps (door opening, food retrieval, predator activation, nest reentry, speed, distance from nest, distance from food). The simplified formula that describes the full GLM model is as follows:

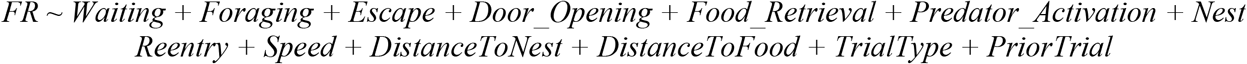

where *FR* is firing rate. The full details of the Lasso GLM model development, validation, and assessment of interaction terms are presented in detail in Amir et al. (2019a) and Kyriazi et al. (2018).

### Statistical Analyses

Data are stated as average ± SEM. In the recording experiments, only cells with stable discharge rates and spike waveforms were considered in the analyses. Statistical tests were two tailed. Different procedures were used to assess statistical significance depending on the data type, as detailed below.

#### Behavior

In the inactivation experiment, we used a mixed-model ANOVA followed by post-hoc paired-samples t-tests to examine the effect of with infusion site (BL, CMT, Control) as between-subject variable and treatment (vehicle, muscimol). In the unit recording experiments, to compare the incidence of a behavior in two conditions, we computed Chi-square tests for independence with a threshold alpha of 0.05. To compare the duration of a behavior in two conditions, we computed a Student’s t-test.

#### Task-related changes in firing rates

To determine whether individual neurons showed statistically significant task-related variations in firing rates, we computed Kruskal-Wallis one-way ANOVAs and applied a Bonferonni correction of α for the number of neurons considered (0.05/716 or 0.000014). For those cells with a significant ANOVA, we then computed Tukey-Kramer post-hoc tests with a threshold α of 0.05. To assess the influence of the predator (presence/absence) and prior trial outcome (success/failure) on firing rates, we computed a Friedman test followed by post-hoc Wilcoxon tests with Bonferroni-corrected α for the number of comparisons.

#### GLM normalized peak firing rate modulations

The absolute peak modulation associated with each variable was normalized to baseline firing rates according to this equation: (Peak-Baseline)/(Peak+Baseline). We kept the sign of the normalized peak to distinguish cells inhibited or excited by each variable. Normalized peak modulations with values ≤0.001 were set to zero because considered non-informative. The average modulation by a task feature was assessed by averaging the absolute value of all peak unit modulations. We assessed the relationship between the modulations associated with the variables across cell types by computing a rank-based (Spearman) correlation. To compare the model fit to observed spiking, we used the coefficient of determination (R^2^) as follows:

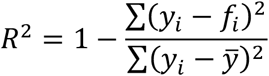

where *y_i_* represents the observed unit spiking and *f_i_* is the model-estimated firing at different time points *i*, and 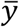 is the overall average of the observed unit spiking.

## RESULTS

### Experiment 1: Muscimol infusions in BLA and CMT

Rats were trained to leave a nest-like compartment to retrieve a food pellet in a brightly lit and elongated arena (**Fig. 1B**). On some trials, rats were confronted with a mechanical predator, which was located at the other end of the foraging arena, facing rats as they approached the food pellet. Trials started when the gateway to the foraging arena opened. After a variable delay, rats moved to the gateway and paused at the door threshold (Waiting phase). Eventually, they left the nest to retrieve a food pellet placed 12.5 to 150 cm from the nest (Foraging phase). Upon retrieving the food pellet (or failing to do so), rats abruptly changed direction and ran to the nest (Escape phase). Upon nest reentry (Nest phase), the gateway was closed. On predator trials, each time rats came within ~25 cm of the food pellet, the predator surged forward, snapped its jaw repeatedly, and returned to its original position.

We compared behavior in the foraging task, 15-30 min after bilaterally infusing the same volume of vehicle (ACSF, 0.1 μl) or fluorescent muscimol (0.9 nanomoles) in BLA or CMT. We formed a third (control) group out of CMT rats with incorrect cannula placements, resulting in the following samples: 10 BLA rats, 8 CMT rats, and 11 control rats. See **figure 1C** for examples of fluorescent muscimol infusions and **figure 2** for schemes illustrating cannula tip locations in the three groups. Vehicle and muscimol infusions were performed on different days, 24 h apart. To minimize the influence of habituation to the predator, the order of the infusions was counterbalanced in all groups.

**Figure 2.**
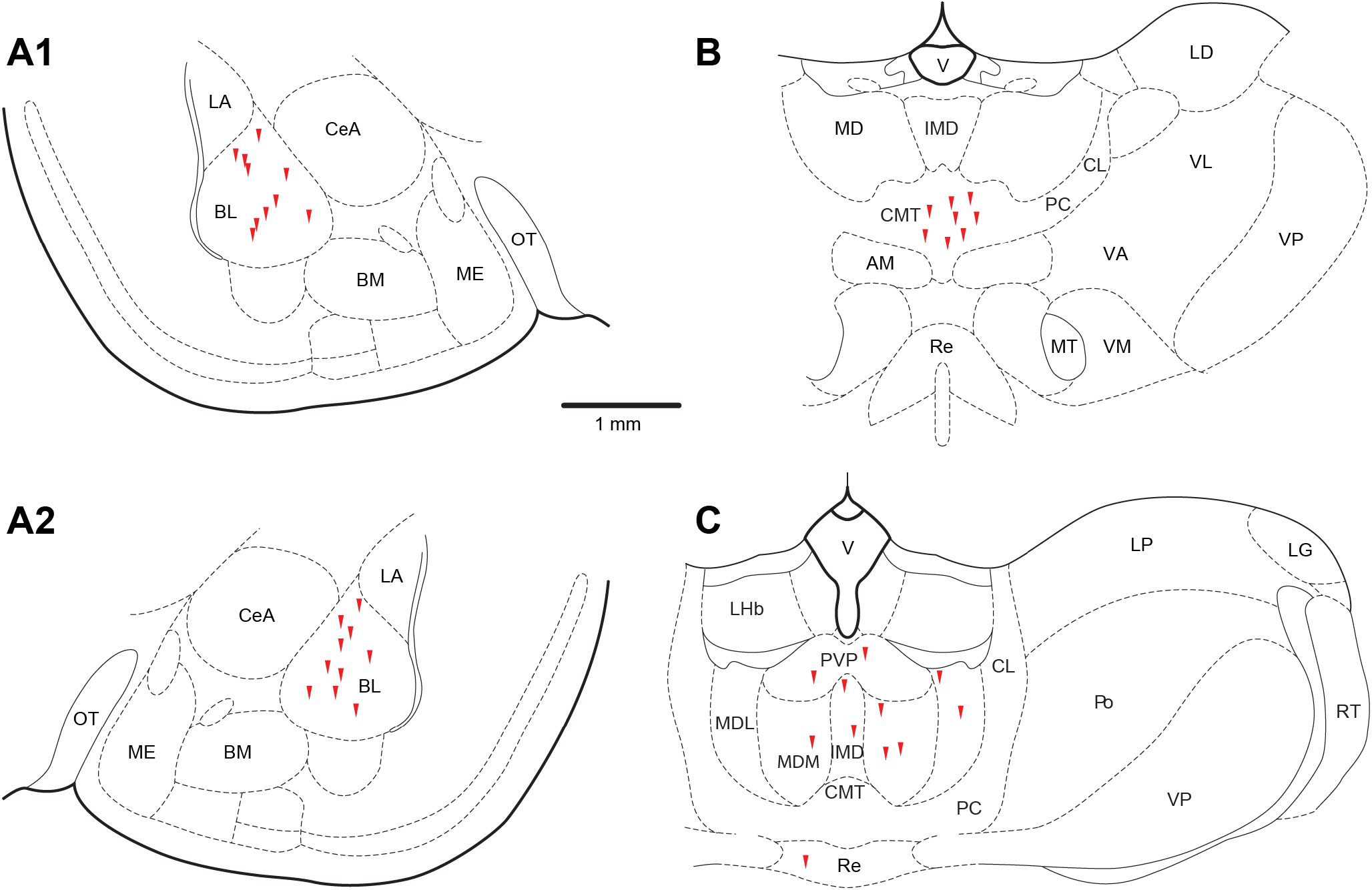
Location of cannula tips (red arrowheads) in the BLA (**A**), CMT (**B**), and control groups (**C**).

Our behavioral protocol adhered closely to that used by Choi and Kim (2010), with one exception: the nest and foraging arena were separated by a sliding door, whose opening signaled the start of a trial. In brief, after habituation to the behavioral apparatus (Days 1-2) and training on the foraging task in the absence of the predator (Days 3-9), rats received a vehicle or drug infusion 15-30 min before giving them the opportunity to retrieve a food pellet placed in the foraging arena, 75 cm from the nest (Days 10-12). When rats advanced within ~25 cm of the food, the predator surged forward (80 cm at 60 cm/s), snapped its jaw repeatedly, and returned to its original position. If the rat successfully retrieved the food pellet, the distance was increased in steps of 25 cm until the rat failed. When the rat failed at 75 cm, the distance was decreased to 50 cm, 25 cm, then 12.5 cm until the rat retrieved the pellet. Trials were separated by 1-2 min intervals during which rats were in the nest with the door closed.

The dependent variables we monitored were the *maximal foraging distance* (**Fig. 1D**) and the amount of time rats hesitated at the door threshold before initiating foraging (hereafter termed ‘*waiting time*’; **Fig. 1E**). For both dependent variables (with Bonferroni-corrected α=0.025), a two-way mixed-model ANOVA with infusion site (BL, CMT, Control) as between-subject variable and treatment (vehicle, muscimol) as within-subject variable revealed a significant effect of infusion site on foraging distance (F(1,26)=5.258, *p*=0.012, η^2^=0.288) but not on waiting time (F(1,26)=0.116, *p*=0.891), a significant effect of treatment (foraging distance: F(1,26)=86.911, *p*<0.001, η^2^=0.770; waiting time: F(1,26)=7.948, *p*=0.009, η^2^=0.234), and a significant interaction between infusion site and treatment on foraging distance (F(226)=11.887, *p*<0.001, η^2^=0.478), but not on waiting time (F(1,26)=2.381, *p*=0.112).

Post-hoc paired-samples t-tests confirmed Choi and Kim’s (2010) observations that muscimol infusions in BL caused a significant increase (320±41%) in maximal foraging distance (t(9)=7.236, *p*<0.001, *d*=2.289). We also found that BL inactivation significantly reduced (−58±19%) waiting times (t(9)=-2.835, *p*=0.020, *d*=0.896). Of the other groups, only CMT infusions were found to have significant effects: a 123% increase in foraging distance (t(7)=8.000, *p*<0.001, *d*=2.829) and a 72±33% decrease in waiting times (t(7)= −2.014, *p*=0.066 *d*=0.768).

### Experiment 2: Unit recordings in CMT during the foraging task

We recorded unit activity in the CMT while rats (n=7) performed the foraging task. **Figure 3A** summarizes the experimental timeline. On each recording day, we conducted alternating trial blocks with (n=10–20) or without (n=10–15) the predator, for a total of 100–120 trials per day. On each trial, the distance between the nest and food pellet was varied randomly between 12.5 and 150 cm. Thus, on predator trials, the proximity of the food pellet to the predator varied randomly on a trial-by-trial basis. Rats showed signs of increased apprehension on predator trials. Specifically, the proportion of aborted trials, that is trials in which after hesitating at the door threshold, rats retreated into the nest instead of initiating foraging, was ~3.5 times more frequent in the presence (2.1%) than absence (0.6%) of the predator (Chi-square test for independence, χ^2^=8.40, 28 sessions from seven rats, *p*=0.003). Moreover, when rats did initiate foraging in the presence of the predator, the retrieval interval increased by 265±34% (Paired-samples t-test; t(27)=6.910, *p*<0.001, *d*=1.306) and the proportion of successful trials dropped 42% (Chi-square test for independence; χ^2^=506.74, 28 sessions from seven rats, p<0.001). Finally, rats spent much longer retrieving the food pellet on predator trials that followed failed (61.3 ± 7.7 s) than successful trials (24.5 ± 3.1 s; Paired-samples t-test; t(26)= –6.084, *p*<0.001, *d*=1.171).

**Figure 3.**
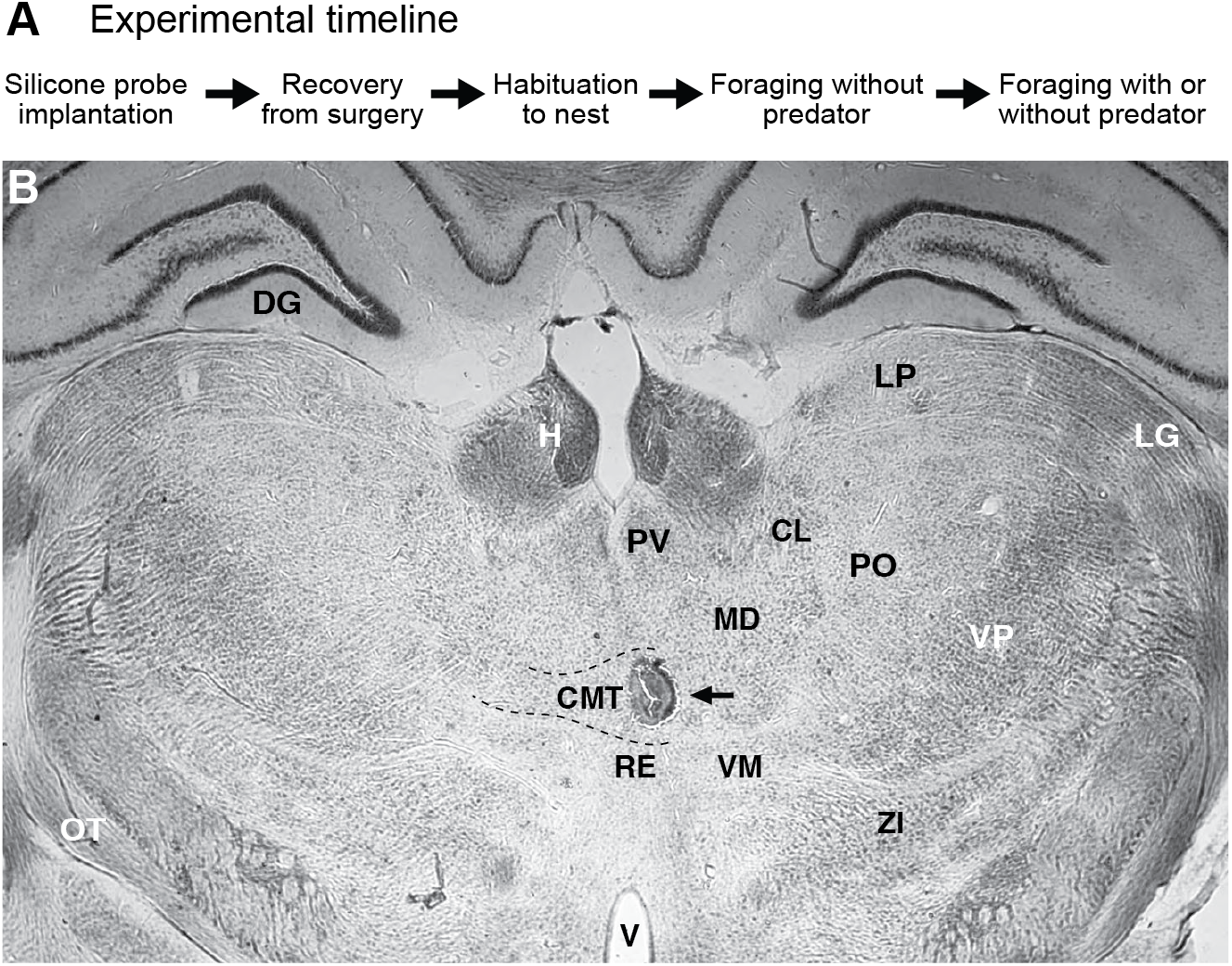
Experimental timeline and histological determination of recording sites. (**A**) Main phases of the experiment. (**B**) Coronal section counterstained with thionine. **Arrow:** electrolytic lesion performed at the conclusion of the experiment to mark the last recording site.

### Task-related activity of CMT neurons

A total of 716 single units, histologically determined to have been located within CMT (**Fig. 3C**), were recorded while rats performed the foraging task. We begin by describing the activity of CMT neurons irrespective of whether the predator was present or absent. We will return to the impact of the predator in a subsequent section.

**Figure 4** shows four representative examples of CMT neurons with obvious task-related activity. For each cell, we provide spike rasters centered on the onset of particular task events and below them, the corresponding average ± SEM firing rate. As the duration of each behavioral phase varied from trial to trial, we rank-ordered the trials based on the timing of events that occurred just before (door opening, left column) or after the behavior of interest (escape, middle column; nest reentry, right column). These events are indicated by blue tick marks. Finally, the cyan lines indicate the average speed of the rats (upward and downward deflections indicate movements away from vs. toward the nest, respectively).

**Figure 4.**
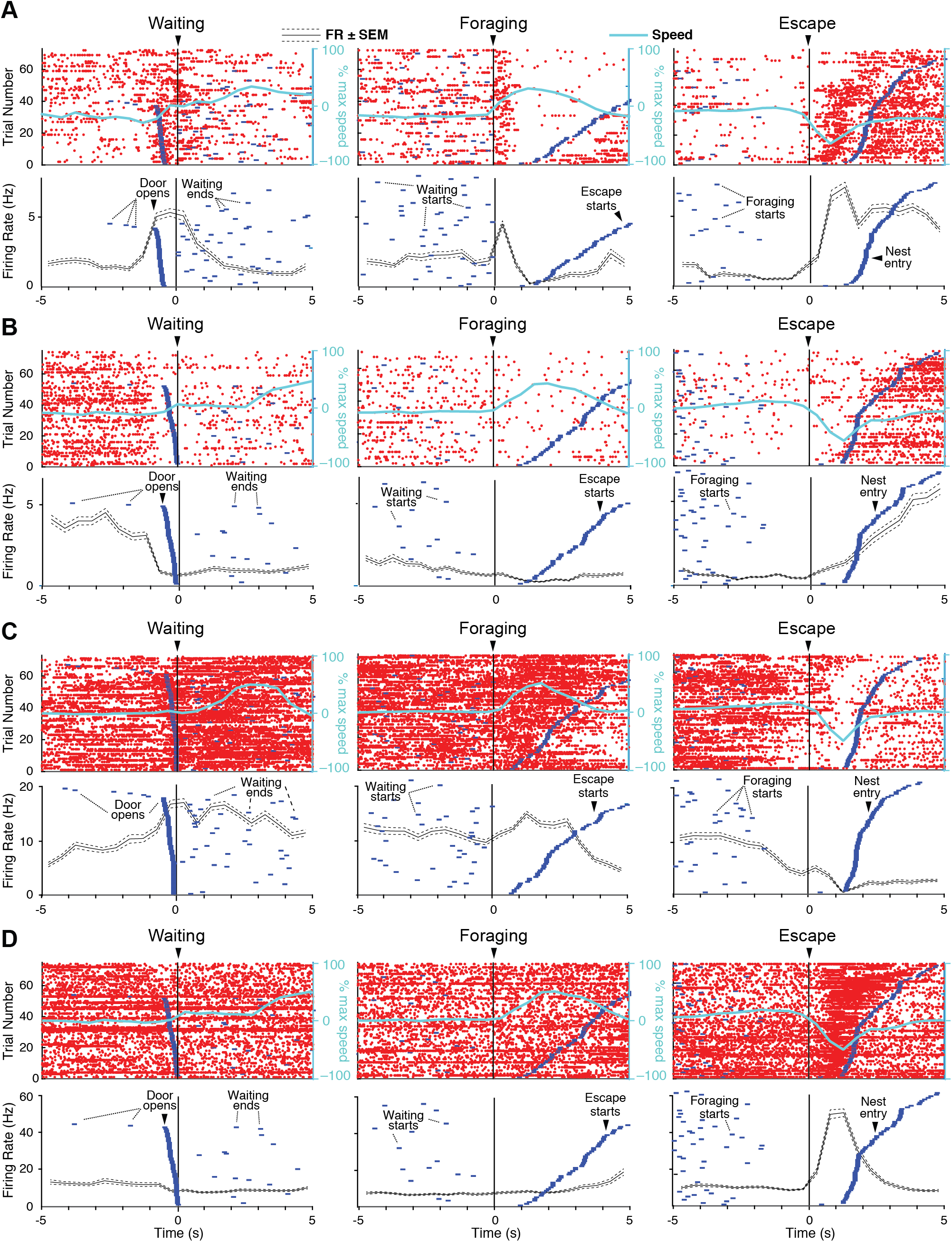
Examples of CMT unit activity during the foraging task. (**A-D**) Activity of four different CMT neurons around the onset of waiting (left column), foraging (middle column), and escape (right column). Top row: rasters where red dots mark spike times and each horizontal line corresponds to a trial. Bottom row: average ± SEM firing rates across all trials depicted in the rasters. In the rasters, trials are rank ordered based on the timing of events that occurred just before (door opening, left column) or after the behavior of interest (escape, middle column; nest entry, right column). These events are indicated by blue tick marks. Cyan lines: rats’ average speed (upward and downward deflections indicate movements away from vs. toward the nest, respectively).

As shown in **figure 4**, the task-related activity of CMT neurons was extremely variable with some cells showing transient increases in firing rates in relation to most task events (**Fig. 4A**), others exhibiting persistent firing rate reductions from the start of the trials until rats returned to the nest (**Fig. 4B**), and others displaying more differentiated activity profiles (**Fig. 4C,D**). To determine whether each neuron showed statistically significant task-related variations in activity, we computed Kruskal-Wallis one-way ANOVAs on firing rates (Bonferonni-corrected threshold α of 0.05/716 or 0.000014) during the baseline (in the nest with the door closed), waiting, foraging, and escape periods. Overall, 88% of CMT cells (633 of 716) showed significant task-related activity (*p*<0.000014). Tukey-Kramer post-hoc tests with a threshold α of 0.05 revealed that 337 of the 633 cells had a significantly different firing rate in at least one of the task phases relative to baseline.

**Table 1** shows the distribution of these 337 cells, specifically how many of them increased or decreased their firing rates relative to baseline in the various task phases. A cursory look at this table reveals that CMT cells are heterogeneous: neurons with increased or decreased activity levels relative to baseline are seen across all task phases. Whereas a higher proportion of cells had increased than decreased firing rates in the waiting, foraging, and nest reentry phases, the opposite was seen for door opening. For escape, our sample was evenly split between increased and decreased activity levels. Of note, many CMT neurons (229 of 337 or 32% of the total 716 neurons) had significantly different firing rates in two or more task phases relative to baseline (**Table 2**).

**Table 1.** CMT cells with statistically significant changes in firing rates in the foraging task. We computed Kruskal-Wallis one-way ANOVAs on the firing rates of CMT neurons (n=716) across the various task phases (bold headings). Of the 633 cells with a significant ANOVA, 337 had firing rates that differed significantly between baseline and one or more task phases (Tukey-Kramer post-hoc test, *p*<0.05). The table lists the number and proportion of significant cells for each task phase. Because many cells had significant changes in firing rates in more than one task phase, the sum of the cells across the different task phases is greater than 337. Abbreviation: FR, firing rate.

|  | Door opening | Waiting | Foraging | Escape | Predator activation | Nest entry |
| --- | --- | --- | --- | --- | --- | --- |
| **CMT (n=716)** | 193<br>(27.0%) | 130<br>(18.2%) | 199<br>(27.8%) | 229<br>(32.0%) | 162<br>(22.6%) | 177<br>(24.7%) |
| **↑ FR** | 69<br>(9.6%) | 86<br>(12.0%) | 117<br>(16.3%) | 113<br>(15.8%) | 89<br>(12.4%) | 100<br>(14.0%) |
| **↓ FR** | 124<br>(17.3%) | 44<br>(6.1%) | 82<br>(11.5%) | 116<br>(16.2%) | 73<br>(10.2%) | 77<br>(10.8%) |

**Table 2.** Number of cells with firing rates significantly different from baseline in one or more task phases. The bold numbers at the top indicate the number of task phases. Same statistical procedures as in the previous table.

|  | Number of Task Phases |  |  |  |  |
| --- | --- | --- | --- | --- | --- |
|  | 1 | 2 | 3 | 4 | 5 |
| **CMT**<br>**(n=716)** | 108<br>(15.1%) | 85<br>(11.9%) | 73<br>(10.2%) | 48<br>(6.7%) | 23<br>(3.2%) |
| **↑ FR** | 49<br>(6.8%) | 46<br>(6.4%) | 46<br>(6.4%) | 24<br>(3.4%) | 9<br>(1.3%) |
| **↓ FR** | 59<br>(8.2%) | 39<br>(5.4%) | 27<br>(3.8%) | 24<br>(3.4%) | 14<br>(2.0%) |

A better appreciation of cell-to-cell variations in task-related activity can be gained by inspecting **figure 5A1**, which shows the activity of all cells with a significant Kruskal-Wallis ANOVA (n=633), z-scored based on variations in firing rates during the baseline period. Cells were rank ordered from least (top, blue) to most active (bottom, yellow) based on their activity during the foraging phase and the same order was kept in the other phases. **Figure 5A2** plots their average (± SEM) firing rates. Comparing the color distributions in the different phases (**Fig. 5A1**) reveals that activity during foraging is a poor predictor of that in other task phases. In fact, rank correlations between z-scored activity in the various phases are consistently low, in the −0.2 to 0.2 range (**Fig. 5B1**). The lack of consistent association between CMT activity in the different phases is highlighted in **figure 5B2**, which plots the ranks of all significant cells in the waiting, foraging, escape, and nest phases, but color coded based on their activity during waiting. Nevertheless, as a group, CMT cells increased their firing rates (0.5–0.7 z score) in all phases of the foraging task until rats returned to their nest (**Fig. 5A2**).

**Figure 5.**
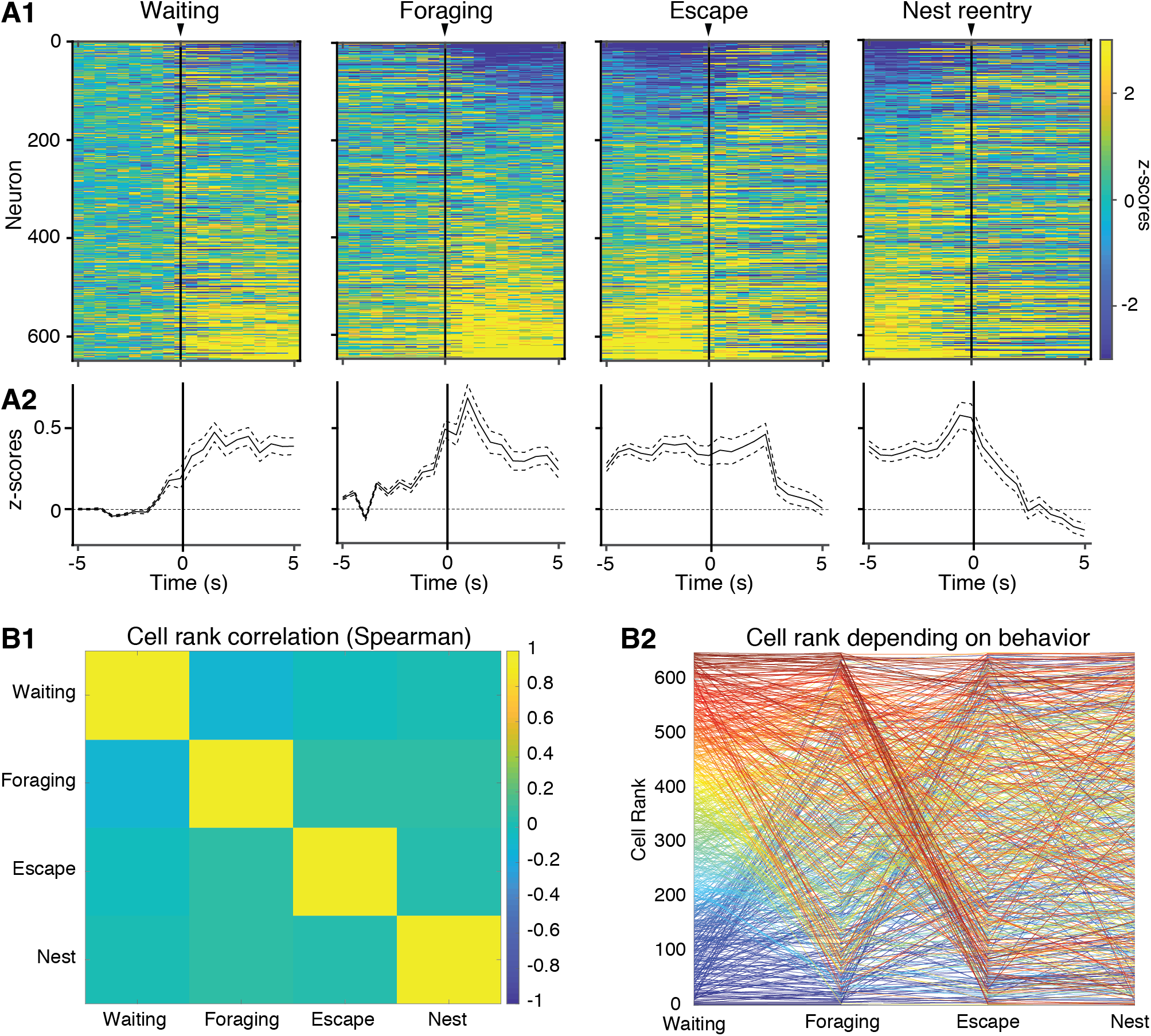
Variations in the activity of CMT neurons during the different phases of the foraging task. (**A1**) Activity of all CMT neurons with significant task-related activity (n=633) during the foraging task. Firing rates were z-scored based on the cells’ activity during the baseline period (in the nest with door closed). Cells were rank-ordered from least (blue) to most active (yellow) based on their activity during the foraging phase and the same order was kept for the other phases. (**A2**) Average ± SEM of z-scored firing rates for all cells in the corresponding panel. (**B1**) Matrix of rank correlations of CMT cells (n=633) between the different task phases. (**B2**) Relationship between rank of same CMT cells in the different task phases. Cells were rank-ordered from least (blue) to the most active (yellow) based on their activity in the Waiting phase. The color-coding of each cell was then kept for the other behavioral phases.

In an attempt to identify subsets of CMT neurons with distinct profiles of task-related activity, we used unsupervised clustering (*kmeans* function in MATLAB). Using raw firing rates, the optimal number of identified clusters by gap statistics was always near the maximum number of tested clusters. Separately averaging the cells’ activity within each cluster failed to reveal distinct patterns of task-related activity (not shown). Similarly inconclusive results were obtained when clustering was carried out on normalized firing rates (z-scored based on variations during the baseline period).

### Influence of the predator and prior trial outcome

As mentioned above, rats showed signs of increased apprehension on trials carried out in the presence of the predator. That is, they waited longer before initiating foraging and foraged more tentatively, especially if they had failed to retrieve the food on the prior trial. To examine the neuronal correlates of these behavioral variations, we compared the z-scored activity of CMT cells on no predator vs. predator trials, rank ordering the cells based on their activity in the foraging phase of no predator trials and keeping the same ordering in the other conditions (**Fig. 6A**). Predator trials were subdivided in two groups, depending on whether rats had failed or succeeded to retrieve the food pellet on the prior trial (failed trials were rare in the absence of the predator). As there were large differences in phase duration across trial types, we plotted the data of **figure 6A** as a function of relative time, by distributing a fixed number of samples evenly across each of the phases of interest.

**Figure 6.**
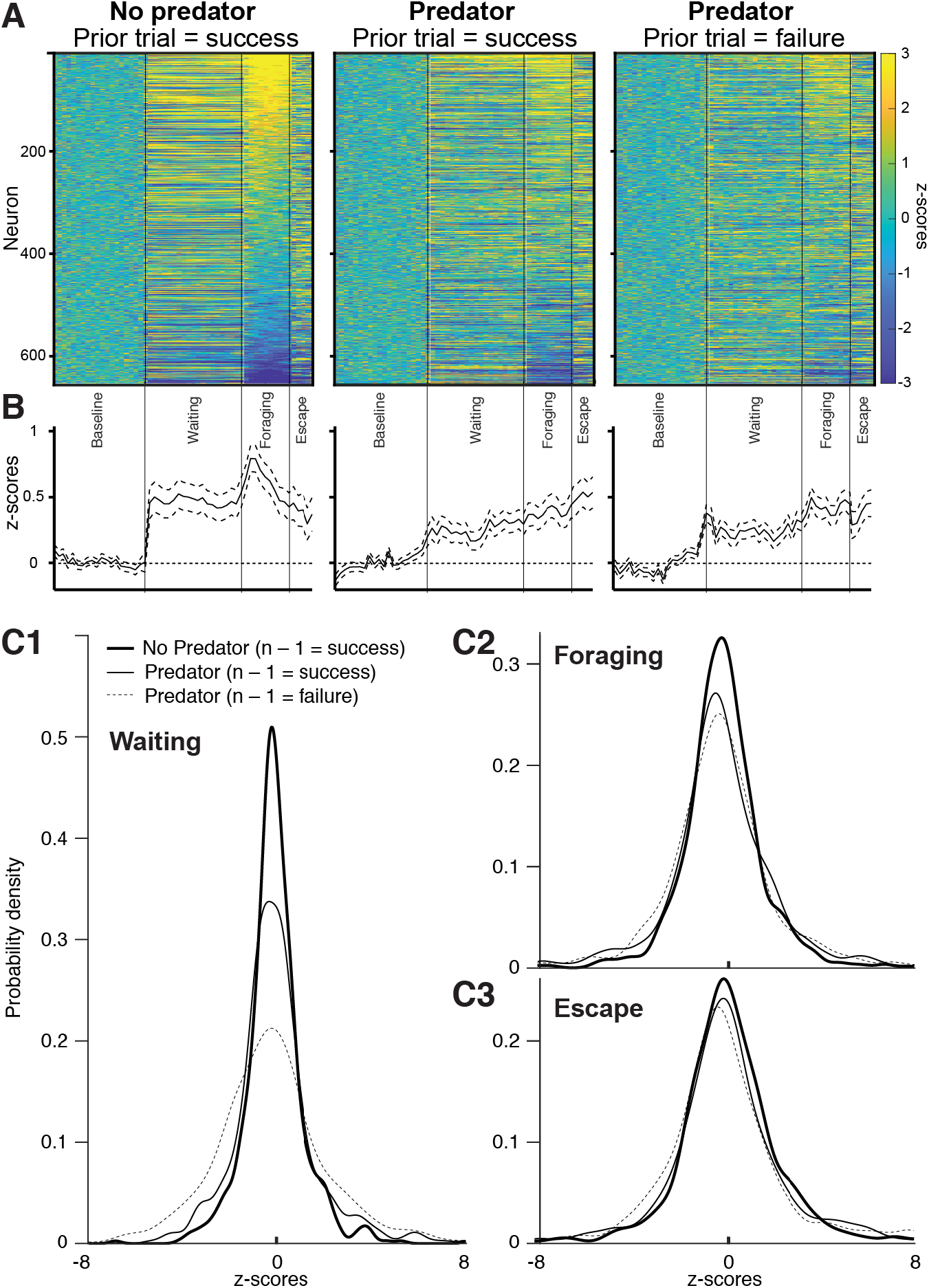
Variations in the activity of CMT neurons as a function of trial type. (**A**) Activity of all CMT neurons with significant task-related activity during the foraging task (n=633) in no predator (left) or predator (middle and right panels) trials. The predator trials were further subdivided as a function of prior trial outcome (success, middle; failure, right). Too few failed trials were available for the no predator condition. In each panel, firing rates were z-scored based on the cells’ activity during the baseline period (in the nest with door closed) and the cells were rank ordered based on their activity on no predator trial during the foraging phase. The same rank order was kept for the other two panels. To allow for comparisons despite trial-to-trial variations in the duration of the different behavioral phases, the data is plotted as a function of normalized time. (**B**) Average of z-scored firing rates ± SEM for all cells in the corresponding panel. (**C**) Kernel density estimation of z-scored and time-relative firing rates in the same trial types as shown in **A** (see legend at top of **C1**) and during different task phases (**C1**, waiting; **C2**, foraging; **C3**, escape).

This analysis revealed that not only were the relative activity levels of CMT cells different across different phases on the same trial type (**Fig. 5A1**), they also differed in the same phase of different trial types (**Fig. 6A1**). Moreover, the average z-scored activity of CMT cells was lower on predator than no predator trials, particularly if rats had failed the prior predator trial (**Fig. 6A2**). A Friedman test confirmed that there was a significant difference in firing rate between different trial types (χ^2^=34.831, df=2, n=195, *p*<0.0001). Post-hoc pairwise comparisons using Wilcoxon tests (with Bonferroni-corrected α=0.017) revealed significantly higher CMT firing rates in *No-Predator* trials compared to the other two trial types (*No-Predator Prior Success* vs. *Predator Prior Success*: Wilcoxon T=5068, Z=3.772, *p*<0.001; *No-Predator Prior Success* vs. *Predator Prior Failure*: Wilcoxon T=5160, Z=4.200, *p*<0.001) but no difference between the two subtypes of predator trials. The difference across trial types can be attributed to changes in the kurtosis of the CMT firing rate distributions as opposed to changes in skewness (**Fig. 6C**). Predator presence and failure in the preceding trial decreased the peakedness of the firing rate distributions.

### Disentangling the correlates of CMT activity

Overall, the above analyses indicate that CMT neurons exhibit large but variable firing rate fluctuations during the foraging task. However, identifying which factors drive these variations in not trivial. Indeed, as foraging trials unfold, a number of temporally overlapping variables fluctuate besides the task phases, such as the rats’ speed and position. To help us quantify the relative influence of these factors, we fit the activity of each CMT unit with a group least absolute shrinkage and selection operator (Lasso) generalized linear model (GLM). This type of GLM exploits variations in the timing and duration of relevant variables to determine the neurons’ encoding preference. Specifically, we considered the rats’ speed, position, distance from the food pellet, the influence of task phases (baseline, door opening, waiting at the door, foraging, predator activation, escape, nest entry), trial types (with or without predator), and prior trial outcome (failure, success). Critically, this type of GLM promotes dimensionality reduction of correlated data and permits sparsity in the identification of the factors linked to neuronal activity (Breheny and Huang 2015; Tibshirani 1996; Yuan and Lin 2006).

**Figure 7** compares the actual firing rates (blue traces) of two CMT cells to that predicted by the model (red traces) for their task variables with the highest beta values. As in these representative examples, the GLM’s output generally matched observed firing rates (see R^2^ values in the upper left corner of each graph). Now at a population level, **Figure 8A** shows frequency distributions of beta values in CMT cells (n=716) for all the variables considered in the GLM (labels at the top left of the histograms), excluding neurons with zero beta values. Excitatory and inhibitory modulations are plotted to the right and left of the histograms’ origin, respectively. For comparison, the same data is provided for a sample of principal BL neurons (n=599) we recorded previously during the foraging task (Amir et al., 2015; **Fig. 8B**). For both cell types, at the top right of each graph, we provide the number of cells with beta values equal to zero, of cells with modulations different from zero, and the average absolute beta for the variable under consideration. In both cell types, note variations in the dispersion of beta values and in the asymmetry of the distributions. For instance, the beta values of CMT cells for ‘door opening’ and ‘nest entry’ show much more variability than for ‘speed’ and ‘distance from food’ (**Fig. 8A**). Yet, the peakedness of CMT distributions (**Fig. 8A**) is generally higher than that of BL distributions (**Fig. 8B**). Moreover, CMT distributions are generally more symmetric than BL distributions, which are typically skewed to the left, betraying a preponderance of negative beta values.

**Figure 7.**
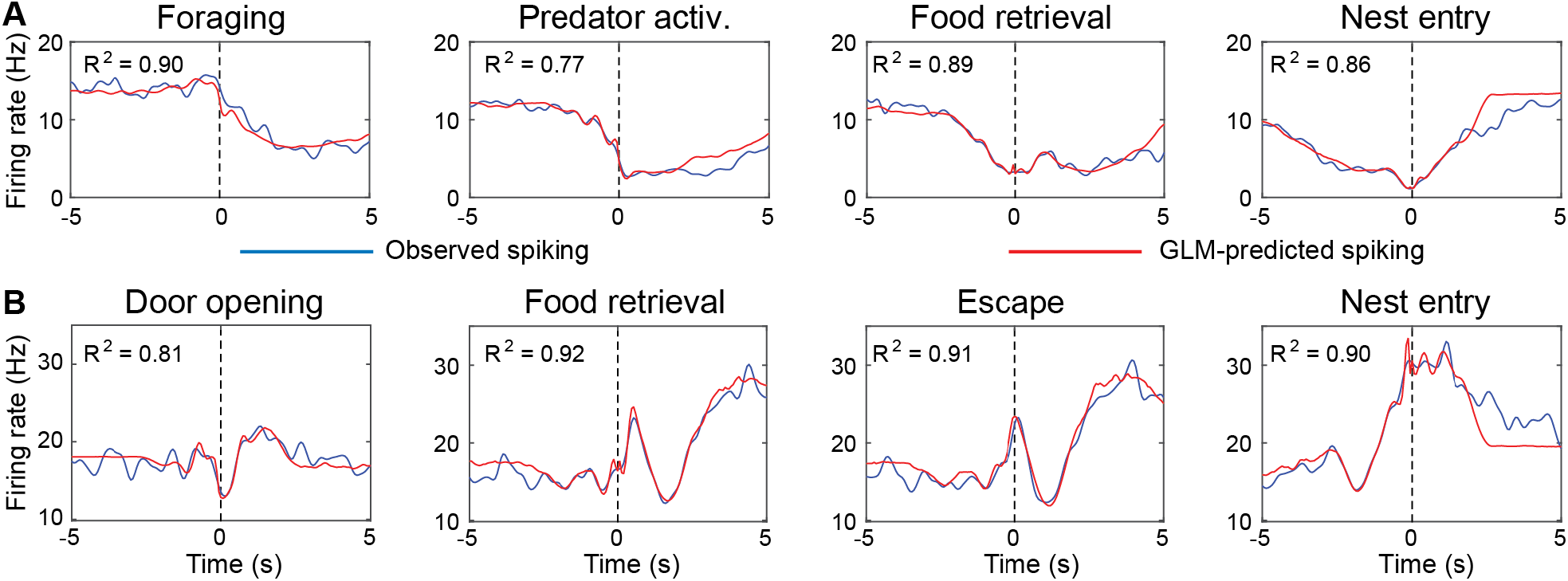
GLM-estimated coding of CMT neurons. **A** and **B** show two different cells. Actual (blue lines) and GLM-predicted (red lines) firing rates during all available trials. In both cells, we show actual and predicted firing rates for the four variables with the largest beta coefficients.

**Figure 8.**
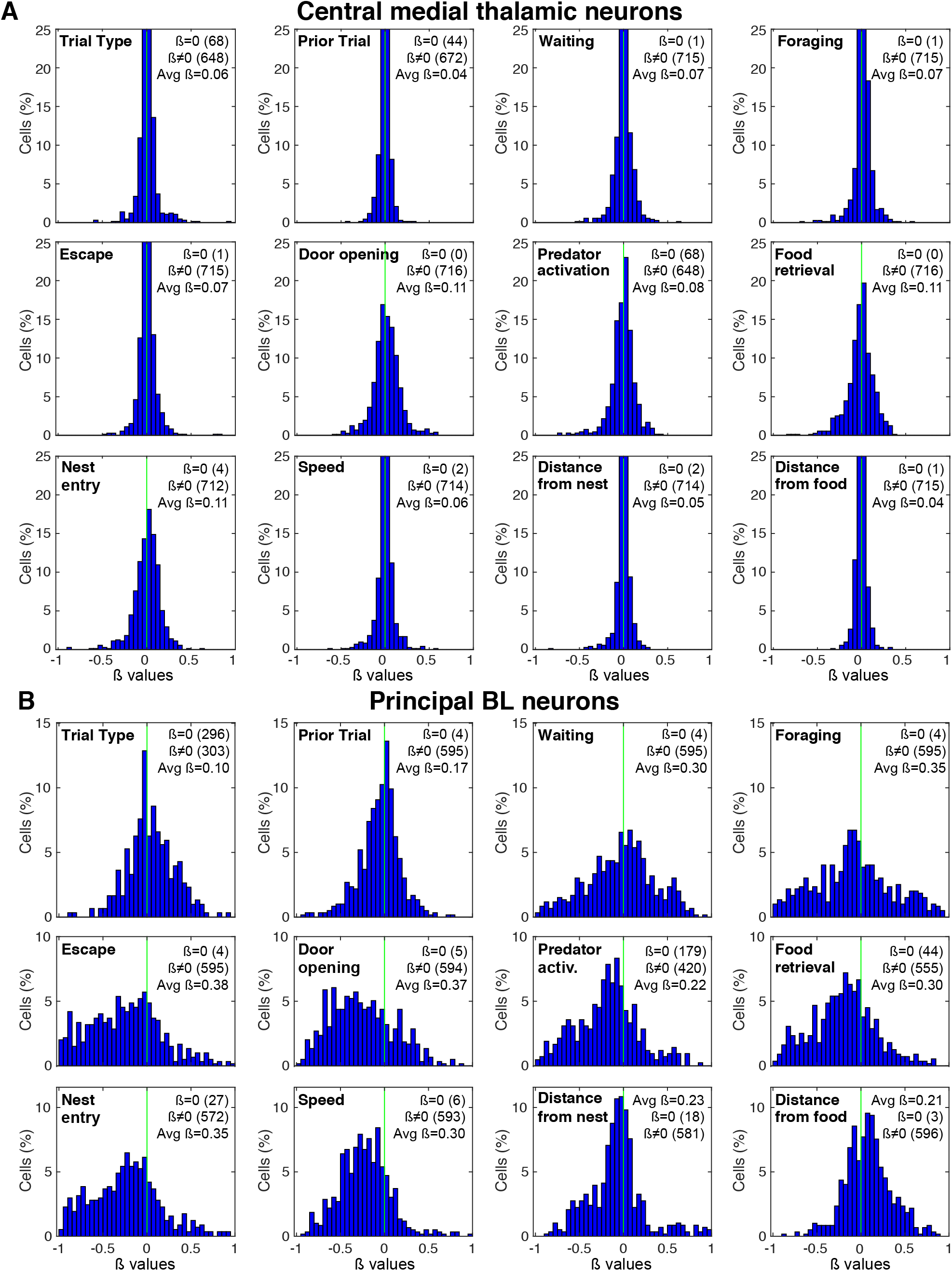
Generalized linear model-estimated coding of 12 variables by CMT and principal BL neurons. (**A**) CMT cells (n=716). (**B**) Principal BL neurons (n=599) recorded in a prior study (Amir et al., 2015). All panels are frequency distribution of firing modulation (*x*-axis; beta values) by the variables indicated at the *top left* of each graph. On the *top right* of each graph, we list (from top to bottom) the number of cells with beta values equal to zero (ß=0), the number of cells with beta values different from zero (ß≠0), and the average absolute beta values (Avg ß).

These contrasting features are also manifest in **figure 9**, which plots for CMT cells (n=716; **Fig. 9A**), principal BL neurons (n=599; **Fig. 9B**), and fast-spiking BL cells (**Fig. 9C**; n=71; also recorded in Amir et al., 2015), their average absolute modulation (**Fig. 9A1, B1, C1**) as well as cumulative excitatory (blue) and inhibitory (red) distributions (**Fig. 9A2, B2, C2**) for all the variables considered in the GLM. Variables were rank ordered from the highest to lowest absolute modulations, in the three cell types separately. Although the variables with the lowest absolute modulations are nearly identical in the three cell types (trial type, prior trial, position, distance from food, speed), there are major differences between the other variables. First, the magnitude of beta values is 2-3 times higher in principal BL neurons than in CMT or fast-spiking BL cells (compare axis range in **Fig. 9B1** vs. **Fig. 9A1, C1**). This difference is likely related to the much higher spontaneous firing rates of CMT and fast-spiking BL neurons compared to principal BL cells (in the nest with door closed: CMT, n=716, 6.09 ± 6.30 Hz; fast-spiking BL cells, n=71, 23.66 ± 13.66 Hz; principal BL cells, n=599, 0.48 ± 1.23 Hz; Kruskal-Wallis ANOVA, df=2, χ^2^=891.65, p<0.0001). As a result, task-related changes in firing rates are proportionally much larger in principal BL cells. Second, whereas inhibitory firing rate modulations are generally higher than excitatory ones in principal BL cells (**Fig. 9B2**), CMT and fast-spiking BL cells show no such imbalance (**Fig. 9A2, C2**). Third, while the variables with the highest modulations were similar in CMT and fast-spiking BL neurons (door opening, food retrieval, nest entry, predator activation), they differed markedly from those dominating the activity of principal BL neurons, particularly escape and foraging.

**Figure 9.**
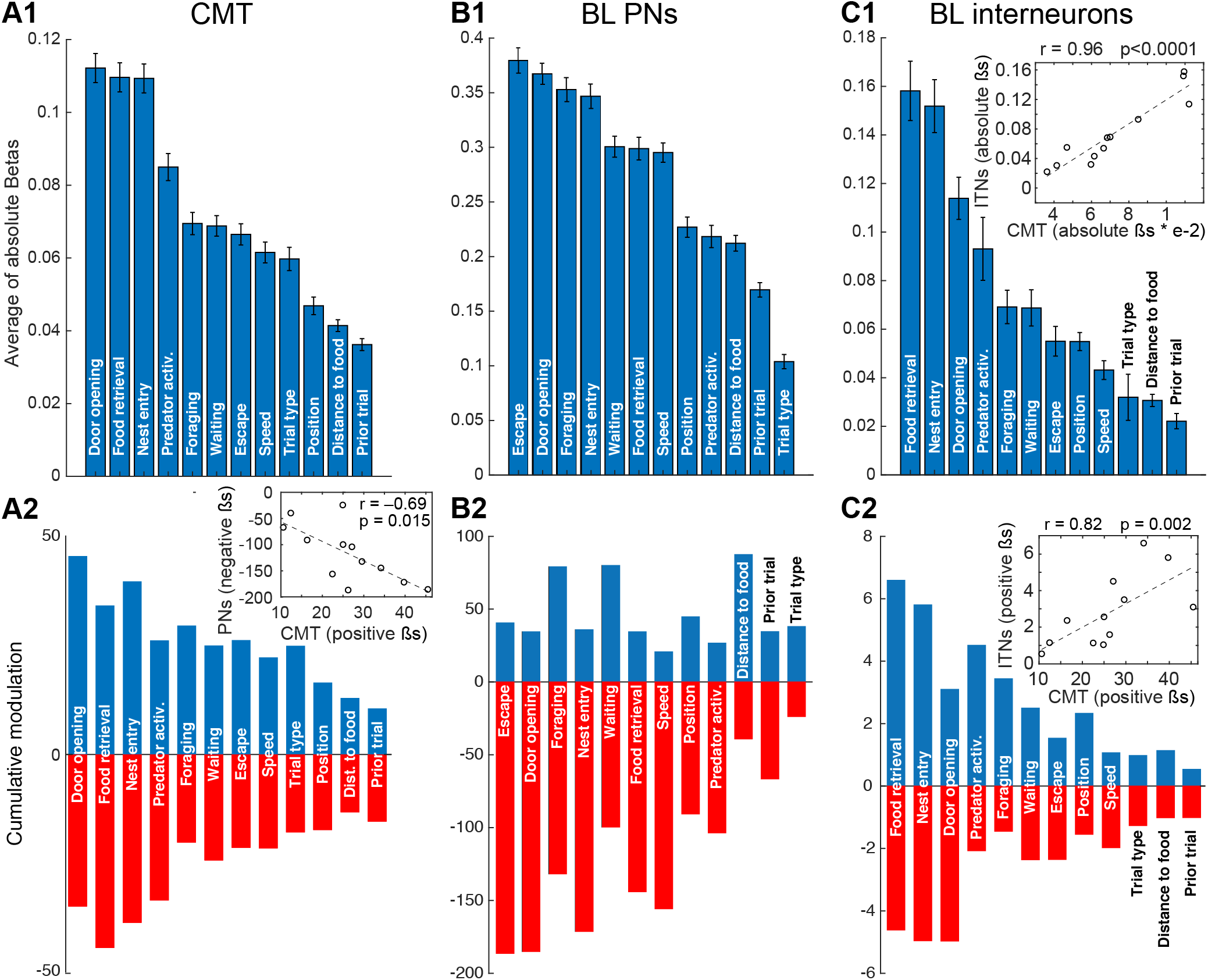
Multidimensional coding by CMT and BL neurons. CMT cells (**A**), principal (PNs, **B**) and fast-spiking (ITNs, **C**) BL neurons recorded in a prior study (Amir et al., 2015). **Panels 1:** average of absolute beta values associated with the 12 different variables considered, rank-ordered by the magnitude of firing rate modulation. **Panels 2:** sum of positive (blue) and negative (red) modulations, ordering the variables as in Panels 1. Inset in **C1** plots average of absolute beta values of BL ITNs (y-axis) vs. that of CMT neurons (x-axis) for the 12 different variables considered. Inset in **A2** plots sum of negative modulation of BL PNs (y-axis) vs. sum of positive modulation of CMT neurons (x-axis) for the same variables. Inset in **C2** plots sum of positive modulation of BL ITNs (y-axis) vs. that of CMT neurons (x-axis) for the same variables. Spearman r’s and associated p values are indicated for each inset.

Consistent with the qualitative similarities between the GLM results obtained in CMT and fast-spiking BL neurons, the average absolute beta values associated with the 12 variables were highly correlated in these two cell types (**Fig. 9C1**, inset; Spearman r=0.96, *p*<0.0001). And so were the positive betas associated with these variables (**Fig. 9C2,** inset; Spearman r=0.82, *p*=0.002). Opposite to this, a negative correlation was found between the positive beta values of CMT cells and the negative beta values of principal BL neurons (**Fig. 9A2**, inset; Spearman r=–0.69, *p*=0.015), raising the possibility that CMT inputs influence BL neurons by activating fast-spiking BL interneurons.

## DISCUSSION

Converging evidence indicates that BL is a major regulator of foraging behavior. In the foraging task for instance, BL inactivation causes rats to become indifferent to the predator (Choi and Kim, 2010). However, at odds with the widely held view that the amygdala detects threats and generate defensive behaviors, unit recordings revealed that most BL neurons have *reduced* firing rates at proximity of the predator (Amir et al., 2015, 2019a). In search of the signals determining this unexpected activity pattern, the present study considered the possible contribution of CMT, which sends a very strong projection to BL (Vertes et al., 2015; Amir et al., 2019b).

In support of the hypothesis that CMT regulates foraging behavior, we observed that inactivation of CMT with muscimol increased foraging distances and decreased waiting times, similar to the effects of BLA inactivation. Moreover, many CMT neurons showed large changes in activity during the foraging task. However, in most task phases, our sample was split between neurons with decreased or increased discharge rates, with a modest bias for the latter. In addition, while GLM analyses revealed that CMT neurons encode many of the same task variables as principal BL cells, the nature and relative importance of the activity modulations seen in CMT neurons differed markedly from those of principal BL cells. By contrast, the GLM revealed striking similarities between the beta values of CMT cells and fast-spiking BL interneurons. Below, we consider the significance of these findings for the regulation of BL activity by CMT neurons.

### CMT neurons have large but heterogeneous activity correlates in the foraging task

Consistent with relay neurons in other thalamic nuclei (Steriade, 1993), CMT neurons fired tonically (median firing rate: ~5 Hz) during quiet wakefulness. In the foraging task, most CMT neurons (88%) displayed significant changes in firing rates in relation to one or more task events. However, in each task phase, the firing rate of many CMT neurons increased while others showed the opposite. Moreover, activity changes developing during any particular task phase were poor predictors of those occurring in other phases.

At a group level, this heterogeneity resulted in modest task related activity. Specifically, average CMT firing rates increased ~0.5 z-scores from baseline to waiting, remained elevated at or slightly above this level (0.7 z-score) through foraging and escape, only returning to, and eventually below, baseline levels after reentry into the nest. Moreover, in the presence of the predator, especially if the rat failed the preceding trial, the overall increase in the firing rates of CMT cells was attenuated. As such, the variability of the predator’s influence (present vs. absent, and success vs. failure) did not impact the heterogeneous activity of CMT neurons, only their overall relative firing rates.

### CMT neurons encode multiple task features

As trials unfold in the foraging task, multiple overlapping variables potentially contribute to alter neuronal activity. These include factors such as the rats’ speed, position in the apparatus, proximity to the food and predator, as well as the rats’ recent experience in the task. Disentangling the relative influence of these multiple factors is especially difficult around the time of predator activation, when rats suddenly switch from foraging to escape. Since peri-event histograms of firing rates cannot dissociate these overlapping factors, we used a GLM, allowing us to assess the influence of each variable while factoring out that of the others by taking advantage of variations in the timing and duration of relevant variables.

As for BL neurons in the foraging (Amir et al., 2019a) and risk-reward interaction tasks (Kyriazi et al., 2018, 2020), this approach revealed that CMT cells encode many task features. That CMT neurons concomitantly represent multiple types of information is consistent with the functionally diverse inputs they receive. Indeed, CMT is the recipient of afferents from the superior colliculus (Krout et al., 2001), parabrachial nucleus (Krout et al., 2000) and many high order cortical areas (Vertes et al., 2015), most of which also project to BL (reviewed in McDonald, 1998).

While the activity of CMT and BL neurons was modulated by a similar array of task features, the relative importance and the nature of these modulations differed. In principal BL neurons, most variables were prevalently associated with inhibitory modulations. This contrasted with CMT and fast-spiking BL interneurons where inhibitory and excitatory modulations were more balanced, with a small preference for excitation.

### Impact of CMT inputs on BL activity in the foraging task

CMT sends a very dense glutamatergic projection to BL (Vertes et al., 2012), which mainly target principal cells (Amir et al., 2019b; Ahmed et al., 2021). However, relative to other extrinsic afferents to the lateral and BL amygdala (LeDoux et al., 1991; Brinley-Reed et al., 1995; Smith et al., 2000; Woodson et al., 2000; Unal et al., 2014), CMT inputs form far fewer synapses with inhibitory interneurons (Amir et al., 2019b), a conclusion supported by a recent physiological study (Ahmed et al., 2021). Based on these results, CMT neurons are not expected to recruit much feed-forward inhibition in BL cells. However, when CMT inputs fire principal BL cells, they strongly recruit feedback interneurons (Ahmed et al., 2021).

Indeed, BL contains the same types of GABAergic neurons as found in cortex (Mascagni and McDonald, 2007; Spampanato et al., 2011; Vereczki et al., 2021). The numerically most important subtype, parvalbumin-immunoreactive (PV^+^) interneurons, many of which have a fast-spiking firing pattern (Woodruff and Sah, 2007), are critically involved in feedback inhibition. They receive strong excitatory inputs from projection cells (Smith et al., 2000; Woodruff and Sah, 2007), they are coupled by gap junctions (Muller et al., 2005; Andrasi et al., 2017), and they form numerous inhibitory synapses on the somatic, axonal, and proximal dendritic domains of projection neurons (Smith et al., 1998; McDonald and Betette, 2001; Andrasi et al., 2017; Veres et al., 2017). Together, these findings raise the possibility that during the foraging task, CMT inputs fire a few principal BL neurons, leading to the recruitment of local feedback interneurons and the consequent inhibition of a high proportion of principal cells.

Two GLM results are consistent with this model. First, we found a nearly perfect correlation between the firing modulations of CMT and fast-spiking BL interneurons (**Fig. 7C**, insets). Second, the modulation of CMT and principal BL neurons were inversely correlated (**Fig. 7A**, inset). Also supporting this model, fluctuations in the population activity of CMT neurons during the foraging task were opposite to that of principal BL neurons. That is, whereas the average firing rate of CMT cells increased during foraging and escape, most principal BL neurons show the converse (Amir et al., 2015, 2019a). Inverse firing rate fluctuations were also observed upon nest reentry, when the firing rate of CMT cells decreased whereas that of principal BL neurons increased (Amir et al., 2015, 2019a). Finally, whereas CMT neurons showed significantly decreased firing rate elevations on predator relative to no predator trials, principal BL neurons showed the opposite (Amir et al., 2015).

### Relation between CMT unit activity and the effects of CMT inactivation

If the above model is correct and the CMT-driven inhibition of principal BL cells is a necessary condition for rats to initiate foraging, how come CMT inactivation results in increased risk-taking? There are two non-exclusive possible explanations. First, besides BL, CMT projects to the dorsal striatum, nucleus accumbens, and multiple cortical areas (anterior cingulate, prelimbic, orbital and insular cortices; Vertes et al., 2012). Thus, it is possible that CMT inactivation affects foraging behavior indirectly, through a disfacilitation of these other sites. Second, since CMT projects to BL and most of CMT’s cortical targets also project to BL (McDonald, 1998), CMT inactivation is expected to cause a major reduction in the excitatory drive to BL neurons. In turn, this “functional deafferentation” may cause a reduction in the firing rate of BL neurons, of comparable magnitude to that seen upon the initiation of foraging.

### Limitations of the foraging task

A limitation of the present study is the use of a pseudo-predator, which imperfectly reproduces the features of an actual predator, particularly with respect to its olfactory cues. Although the pseudo-predator does not fully capture the presentation of natural threats, it did alter the rats’ behavior as would be expected from an actual predator. That is, rats showed signs of increased apprehension on predator trials: they waited longer before initiating foraging, failed to retrieve the food on a higher proportion of trials, and foraged more tentatively, especially if they had failed the prior trial. Critically, we could detect changes in the activity of CMT neurons in relation to these behavioral variations. Hence, it appears that the foraging task captures essential features of prey-predator interactions, justifying its use here and the interpretation of the results.

### Relation to previous work on the regulation of motivated behaviors by thalamic inputs to the amygdala

The role of the thalamus in motivated behavior has received comparatively little attention, possibly because it is thought to support general functions like arousal or the transfer of sensory information (Bradfield et al. 2013; James et al., 2021). However, rapidly accumulating data implicates the mediodorsal (Mair et al., 2021), intralaminar (Bradfield et al., 2013; Bradfield and Balleine, 2017; Cover and Mathur, 2021), and paraventricular (PVT; Do-Monte et al. 2016; Choi and McNally, 2017; Petrovich, 2021) thalamic nuclei in a variety of motivated behaviors. While the contribution of these thalamic nuclei to motivated behaviors are generally thought to depend on their projections to cortex (Saalman, 2014) or striatum (Bradfield et al., 2013; Smith et al., 2014; Bradfield and Balleine, 2017), PVT projections to the amygdala play an important role, particularly with respect to conditioned fear responses. Indeed, inactivation of PVT neurons attenuate conditioned fear responses (Padilla-Coreano et al. 2012; Do-Monte et al. 2015; Penzo et al., 2014, 2015), an effect mediated by PVT projections to the central lateral nucleus of the amygdala (Penzo et al., 2015). Combined with our results, these findings raise the possibility that multiple thalamic inputs regulate foraging behavior via projections to different amygdala nuclei.

## Acknowledgements

This material is based upon work supported by NIH grant R01 MH107239 to Denis Paré.

## Competing financial interest

The authors declare that they have no competing financial interests.

## Notes

### Competing Interest Statement

The authors have declared no competing interest.

